# Experimental Evolution Across Different Thermal Regimes Yields Genetic Divergence in Recombination Fraction But No Divergence in Temperature-Associated Plastic Recombination

**DOI:** 10.1101/238931

**Authors:** Kathryn P. Kohl, Nadia D. Singh

**Affiliations:** Department of Biology, Winthrop University; Department of Biology, University of Oregon

**Author notes:** To whom correspondence should be addressed: Department of Biology, University of Oregon, 5289 University of Oregon, Eugene OR 97403. Data archiving: These data will be submitted to Dryad upon the publication of this manuscript.

**Keywords:** Recombination, phenotypic plasticity, experimental evolution, variable environment

## Abstract

Phenotypic plasticity is pervasive in nature. One mechanism underlying the evolution and maintenance of such plasticity is environmental heterogeneity. Indeed, theory indicates that both spatial and temporal variation in the environment should favor the evolution of phenotypic plasticity under a variety of conditions. Cyclical environmental conditions have also been shown to yield evolved increases in recombination frequency. Here were use a panel of replicated experimental evolution populations of *D. melanogaster* to test whether variable environments favor enhanced plasticity in recombination rate and/or increased recombination rate in response to temperature. In contrast to expectation, we find no evidence for either enhanced plasticity in recombination or increased rates of recombination in the variable environment lines. Our data confirm a role of temperature in mediating recombination fraction in *D. melanogaster*, and indicate that recombination is genetically and plastically depressed under lower temperatures. Our data further suggest that the genetic architectures underlying plastic recombination and population-level variation in recombination rate are likely to be distinct.

## Introduction

From seasonal color variation in butterflies (e.g. Hazel 2002) to nutrient-dependent horn dimorphism in dung beetles (e.g. Emlen 1994), phenotypic plasticity abounds in nature. Though there is little debate regarding its ubiquity and its central role in generating phenotypic diversity, what remains unresolved is twofold: the role of phenotypic plasticity in evolution and how phenotypic plasticity itself evolves (Via et al. 1995; West-Eberhard 2003; de Jong 2005; Pfennig et al. 2010). The genetic and molecular mechanisms mediating phenotypic plasticity lie at the heart of these debates and yet are largely unknown. For instance, it remains controversial whether there are independent ‘plasticity’ genes or whether plasticity in a trait is governed by the same genes that underlie population-level variation in that trait (for review see Sarkar 2004). Much work is thus required to determine the genetic and molecular underpinnings of phenotypic plasticity. However, an understanding of the genetic architecture and molecular basis of phenotypic plasticity is clearly necessary for modeling the evolution of phenotypic plasticity and for determining how phenotypic plasticity may contribute to evolutionary diversification, speciation, and adaptation.

A model trait to address these fundamental questions regarding the genetic architecture of plasticity would satisfy two requirements. One, that trait must exhibit phenotypic plasticity in response to environmental or developmental conditions. Two, that trait would vary genetically as well, which would enable disentangling the genetic basis of population-level variation in the trait from the genetic basis of phenotypic plasticity in that trait. Meiotic recombination rate meets both of these requirements, making it an ideal trait for investigating the genetic basis of phenotypic plasticity.

Meiotic recombination rate is a prototypical example of a trait capable of phenotypic plasticity in many taxa. For instance, social stress has been associated with increased recombination rates in mice (Belyaev and Borodin 1982), and temperature is known to affect rates of crossing-over in Drosophila (Plough 1917, 1921; Stern 1926; Smith 1936; Grushko et al. 1991). Similarly, exposure to parasites has been associated with elevated recombination rates (Andronic 2012; Singh et al. 2015), and nutrient stress is associated with increased recombination rates in yeast (Abdullah and Borts 2001) and Drosophila (Neel 1941). Further, a clear link between maternal age and recombination rate has been found in humans (e.g. Kong et al. 2004; Hussin et al. 2011), mice (Henderson and Edwards 1968; Luthardt et al. 1973) and Drosophila (Stern 1926; Bridges 1927; Redfield 1964; Lake 1984; Chadov et al. 2000; Priest et al. 2007; Tedman-Aucoin and Agrawal 2012; Hunter et al. 2016b). Thus, meiotic recombination rate shows variability in the context of organismal development and across environments, making it a model trait for exploring the evolution of and genetic architecture of phenotypic plasticity.

Meiotic recombination rate itself is also genetically variable in a variety of systems including Drosophila (e.g. Brooks and Marks 1986; Hunter et al. 2016a), humans (e.g. Fledel-Alon et al. 2011), mice (e.g. Dumont et al. 2009), and other mammals (e.g. Sandor et al. 2012; Ma et al. 2015; Johnston et al. 2016). In mammals, several genes have been identified that have been consistently linked to population-level variation in recombination rate in multiple species (Kong et al. 2008; Chowdhury et al. 2009; Hinch et al. 2011; Sandor et al. 2012; Capilla et al. 2014; Johnston et al. 2016). The genetic architecture of recombination rate in Drosophila appears to be distinct, however (Hunter et al. 2016a), and population-level variation in *D. melanogaster* appears to be governed by a large number of loci each of which has a small effect on recombination.

Here we exploit an experimental evolution framework to explore the genetics and evolution of recombination and plastic recombination in *D. melanogaster*. Specifically, we used a panel of replicated experimental evolution populations of *D. melanogaster* (Yeaman et al. 2010). These populations were evolved in either a constant environment (16°C or 25°C) or in a fluctuating environment in which individuals were alternated between 16°C and 25°C every generation. Because recombination is sensitive to temperature in Drosophila, we leveraged this panel to determine how recombination rate and plastic recombination evolve in response to these experimental regimes.

To generate hypotheses that can be tested within this experimental evolution context, we consider the theoretical framework surrounding the evolution of plastic recombination and the evolution of increased recombination. What conditions favor the evolution of plastic recombination? Theory has shown quite clearly that plastic recombination readily evolves in haploid systems if recombination is fitness-dependent (Hadany and Beker 2003). That is, modifiers that facilitate recombination in poor quality individuals but prevent recombination in high quality individuals can successfully invade populations under a large range of conditions. Fitness-dependent recombination is thus a solution through which the benefits of recombination (bringing together favorable combinations of alleles) can be realized without suffering the consequences of recombination (breaking apart favorable combinations of alleles). However, fitness-associated recombination evolves less readily in diploids (Agrawal et al. 2005), at least as a consequence of the direct effects of the recombination modifier (Agrawal et al. 2005; Rybnikov et al. 2017). However, fitness associated recombination may evolve in diploids as a consequence of selection on average recombination rate if there is cis-trans epistasis or if there are maternal effects on fitness (Agrawal et al. 2005). Moreover, recent work has showed that in a cyclical, two state-environment, condition-dependent recombination strategies are favored over constant recombination strategies in a diversity of circumstances, as a consequence of both direct and indirect effects of the plastic modifier (Rybnikov et al. 2017).

With respect to the evolution of recombination, theory predicts that in a diploid system, modifiers that increase recombination may invade populations under certain conditions. If recombination is favored because it heightens the efficacy of directional selection, then increased recombination can evolve if epistasis is weak and negative (Barton 1995; Otto and Michalakis 1998). Increased recombination can also evolve in fluctuating environments, wherein combinations of alleles that are beneficial in the current environment become deleterious in a future environment (Charlesworth 1976; Otto and Michalakis 1998; Dapper and Payseur 2017). However, the periodicity of environmental change is a critical determinant of the evolutionary trajectory of recombination rate modifiers. If the sign of linkage disequilibrium varies cyclically and the period of fluctuations is greater than two, then increased recombination may evolve (Charlesworth 1976; Carja et al. 2014). Environmental fluctuations that occur over longer time scales have similar evolutionary dynamics to the directional selection scenario, in which weak negative epistasis between alleles favored in the current environment is required for the evolution of increased recombination (Barton 1995; Otto and Michalakis 1998).

Theory thus predicts that cyclical environments can affect both the evolution of recombination and the evolution of plastic recombination. Given that cyclical two-state environments favor condition-dependent recombination over constant recombination, we hypothesized that the populations in the variable thermal regime would evolve a greater magnitude in temperature-associated plastic recombination relative to those populations evolving under a constant temperature. We further hypothesized that flies in our fluctuating environment experimental regime would evolve increased overall rates of recombination relative to their counterparts evolving under constant temperature conditions. In contrast to this expectation, we find no evidence for increased plastic recombination in the fluctuating environment selection lines. In addition, our prediction of increased recombination in the fluctuating temperature regime is not borne out by our data; these populations instead exhibit recombination rates intermediate between the two constant-temperature evolved populations. Our data also confirm a role of temperature in mediating recombination fraction in *D. melanogaster*. Interestingly, our data indicate that recombination is genetically and plastically depressed under lower temperatures, independent of experimental evolution treatment. We find significant differences in recombination frequency between the two constant temperature treatments at both experimental temperatures, indicating that recombination rate has evolved over the course of the experimental evolution regime through direct or indirect selection. These observations collectively suggest that the genetic basis of plastic recombination is independent from the genetic basis of population-level variation in recombination fraction.

## Materials and Methods

### Experimentally evolved populations

The experimentally evolved populations used in the present study were generated previously and are described in detail elsewhere (Yeaman et al. 2010; Cooper et al. 2012; Condon et al. 2014). Briefly, wild-caught *Drosophila melanogaster* females from British Columbia were used to establish 298 isofemale lines. Progeny from these isofemales lines were used to establish a large breeding population that was allowed to grow for six generations. This breeding population ultimately reached a population size of ~64,000 adults. This population was maintained for nine generations and was subsequently used to found the experimental evolution populations.

There were five replicate populations for each of three experimental treatments: 1) 16° C constant (C regime), 2) 25° C constant (H regime) and 3) a fluctuating thermal regime in which the flies were alternated between 16° and 25° C every four weeks (T regime). Generation time varied among treatments; a new generation was established at 2-week intervals for flies at 25° C and at 4-week intervals for flies at 16° C. These selective environments were maintained for over three years for a total of 32 generations at 16° C, 64 generations at 25° C, and an intermediate number of generations for the thermally variable experimental evolution regime.

Following this experimental evolution, genotypes were sampled from each of the five replicate populations for each of the three experimental treatments. For two consecutive generations, a single virgin female was mated to a single virgin male to establish an isofemale line. Multiple isofemale lines were established for each of the replicate populations through two generations of brother-sister mating. These lines were then transferred every three weeks for 27 months under controlled conditions. The goals of establishing isofemale lines were to isogenize the genome within each line and to minimize further evolution in the laboratory.

Brandon Cooper generously provided these lines to us in 2012. We maintained these lines in a 12 hour:12 hour light:dark cycle on a standardized cornmeal-yeast-molasses medium. Adult flies were placed at 16° C and 25° C for experimentation, and offspring of these flies were collected for use in the first cross (described below).

### Estimating recombination

To assay recombination rate, we took advantage of visible, recessive markers in *D. melanogaster*. To measure recombination rates on the *3R* chromosome, we used a strain marked with *ebony* (*e^4^*) and *rough* (*ro^1^*); these markers are 20.4 cM apart (Lindsley and Grell 1967). These markers have been used extensively in our lab to estimate recombination frequency (Jackson et al. 2015; Singh et al. 2015; Hunter et al. 2016a; Hunter et al. 2016b).

To assay recombination rate variation in the experimental evolution lines, we used a classic two-step backcrossing scheme. All crosses were executed at either 25° C or 16° C with a 12:12 hour light:dark cycle on standard media using virgin females aged roughly 24 hours. We conducted 1-9 (average 4.5) replicate assays for each line at each temperature. The number of lines assayed from each population at each temperature is presented in Table 1. For the first cross, ten virgin females from each experimental evolution line were crossed to ten *e ro* males in vials. Males and females were allowed to mate for five days, after which all adults were cleared from the vials. F_1_ females resulting from this cross are doubly heterozygous; these females are the individuals in which recombination is occurring. To uncover these recombination events we backcross F_1_ females to doubly-marked males. For this second cross, ten heterozygous virgin females were collected and backcrossed to ten doubly-marked males. Males and females were allowed to mate for five days, after which all adults were cleared from the vials. After eighteen days, BC_1_ progeny were collected and scored for sex and for visible phenotypes. Recombinant progeny were then identified as having only one visible marker (*e*+ or +*ro*). For each replicate, recombination rates were estimated by taking the ratio of recombinant progeny to the total number of progeny. Double crossovers cannot be recovered with this assay, so our estimates of recombination frequency are likely to be biased downwards slightly.

**Table 1:** Number of replicate lines assayed in each population at each temperature

| Population | 16°C | 25°C |
| --- | --- | --- |
| C1 | 14 | 14 |
| C2 | 8 | 8 |
| C3 | 13 | 13 |
| C4 | 10 | 10 |
| C5 | 12 | 13 |
| H1 | 11 | 11 |
| H2 | 9 | 9 |
| H3 | 12 | 12 |
| H4 | 10 | 9 |
| H5 | 9 | 9 |
| T1 | 9 | 9 |
| T2 | 11 | 11 |
| T3 | 9 | 9 |
| T4 | 10 | 10 |
| T5 | 12 | 12 |
| **TOTAL** | **159** | **158** |

### Statistical analyses

All statistics were conducted using JMPPro v13.0. We used a generalized linear model with a binomial distribution and logit link function on the proportion of progeny that is recombinant. We treated each offspring as a realization of a binomial process (either recombinant or nonrecombinant), summarized the data for a given vial by the number of recombinants and the number of trials (total number of progeny per vial), and tested for an effect of line, temperature, replicate population, and experimental evolution regime. The lines that had missing data at one of the two temperatures were excluded from the analysis. Note that replicate population is nested within experimental evolution regime and line is nested within replicate population. The full model is as follows:

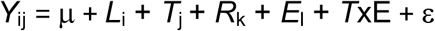

for: *i* = 1…145, *j* = 1…2, *k* = 1…5, *l* = 1…3,

*Y* represents the proportion of progeny that is recombinant, μ represents the mean of regression, and ε represents the error. *L* denotes strain, *T* denotes temperature, *R* indicates the replicate population, *E* represents the experimental evolution regime, and *TxE* denotes the interaction of temperature and experimental evolution regime. All of these are modeled as fixed effects.

We also tested specifically for genetic variation in recombination plasticity using a similar statistical approach. We estimated recombination plasticity as the change in recombination fraction between pairs of replicates at 25 and 16 degrees C, where each member of the pair was randomly chosen (without replacement) from the total number of replicates of that line surveyed at that temperature. If the line had differing numbers of replicates measured at each temperature, the number of replicate pairs used to estimate the change in recombination was limited by the temperature at which fewer replicates were assayed. The extra replicates at the other temperature were not included in the analysis. We tested for an effect of line, replicate population, and experimental evolution regime, where replicate population is nested within experimental evolution regime and line is nested within replicate population. The full model is as follows:

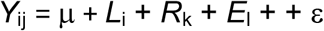

for: *i* = 1…145, *k* = 1…5, *l* = 1…3,

*Y* represents the change in recombination fraction (RF_25_- RF_16_), μ represents the mean of regression, and ε represents the error. *L* denotes strain, *R* indicates the replicate population and *E* represents the experimental evolution regime. All of these are modeled as fixed effects.

Finally, we tested for an effect of temperature on the potential viability effects associated with the doubly marked chromosome. We used a generalized linear model with a binomial distribution and logit link function on the proportion of progeny that is recombinant. We treated each non-recombinant offspring as a realization of a binomial process (either wild-type or *e ro*), summarized the data for a given vial by the number of wild-type flies and the number of trials (total number of non-recombinant progeny per vial), and tested for an effect of line, temperature, replicate population, and experimental evolution regime. Note that replicate population is nested within experimental evolution regime and line is nested within replicate population. The full model is as follows:

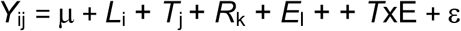

for: *i* = 1…145, *j* = 1…2, *k* = 1…5, *l* = 1…3,

*Y* represents the proportion of non-recombinant progeny that is wild-type, μ represents the mean of regression, and ε represents the error. *L* denotes strain, *T* denotes temperature, *R* indicates the replicate population, *E* represents the experimental evolution regime, and *TxE* denotes the interaction of temperature and experimental evolution regime. All of these are modeled as fixed effects.

## Results

### Robustness of recombination fraction estimation

In total, 149,326 progeny were collected from the experimental crosses and scored for recombinant phenotypes. A total of 65,340 of those progeny resulted from crosses at 16°C while the remaining 83,986 flies resulted from crosses at 25°C. The number of progeny per replicate vial ranged from 10-247, with a mean of 92 progeny per replicate vial at 16°C and 118 flies per vial at 25°C.

To test for deviations from expected ratios of phenotype classes, we performed *G*-tests for goodness of fit for all crosses for the following ratios: males versus females, wild-type flies versus *e ro* flies and finally, *e* + flies versus + *ro* flies. The null hypothesis for each comparison is a 1:1 ratio of phenotype classes. For each of the crosses, we summed progeny counts across all replicates of that cross.

Comparing total females to total males, 16 of 317 (5%) lines show significant deviations from the expected 1:1 ratio (*P* < 0.05, G-test). A relative excess of females is observed in 14 of 16 of those lines. The female-biased lines show M/F ratios ranging from 0.21-0.71, with an average of 0.58 and a median of 0.62, indicating an approximately symmetrical distribution. Of the 16 lines, nine show a bias at 16 degrees, and seven show a bias at 25 degrees. None of the lines show a significant bias at both temperatures. One of these deviations remains significant after using a Bonferroni-correction for multiple tests (Bonferroni-corrected *P* = 0.02, G-test). While the Bonferroni correction is very conservative, we further note that the number of significant tests we observe is not outside the 95^th^ percentile of a binomial distribution with *P* = 0.05; with this *P*-value we expect to see 22 significant tests and we only observe 16.

With respect to wild-type versus *e ro* flies, 30 of 317 (9%, slightly above the 22 tests expected to be positive given binomial sampling) lines show a significant deviation from the expected 1:1 ratio (*P* < 0.05, G-test), and in all but five of these cases these crosses yield a relative excess of wild-type flies. The ratio of wild-type to double mutant in those wild-type biased lines ranges from 1.4-3.8, with an average of 2.0. This is consistent with a mild viability defect associated with the visible markers. Eight of the 30 strains showing deviations exhibit this skew at 16 degrees, with the remaining 22 showing deviations at 25 degrees. After correcting for multiple testing, three H lines maintain a significant deviation from the 1:1 ratio of phenotype classes (Bonferroni-corrected *P* < 0.0005, G-test).

Finally, 19 of 317 (6%, fewer than the 22 positive tests given with binomial sampling) lines show significantly different numbers of *e* + flies versus + *ro* flies (*P* < 0.05, G-test), with 13 of those 19 lines showing an excess of + *ro* flies. The mean and median *e* +/+ *ro* ratio for these 13 lines are 0.34 and 0.35, respectively. Nine of the 19 skewed lines show a skew at 16 degrees while 10 show a skew at 25 degrees. None of these deviations remain significant after using a Bonferroni-correction for multiple tests.

Although the number of lines with significant deviations from null expectation is quite small relative to the total number of lines, especially in the context of binomial sampling, our data are nonetheless indicative of a mild viability defect associated with our marked chromosome. However, these skewed ratios do not appear to depend on temperature as is evidenced by the observation that skewed ratios were observed nearly equally between the two experimental temperatures. Moreover, fitting a generalized linear model with a binomial distribution and logit link function on the proportion of non-recombinant progeny that is wild-type shows no significant effect of temperature (*P* = 0.07, χ^2^ test). We thus believe that whatever small viability defects are associated with the doubly marked chromosome are not systematically biasing the estimates of recombination in this experiment.

### Factors affecting recombination fraction

To identify the factors contributing to the observed variation in the recombination fraction in the current experiment, we used a logistic regression model. We were particularly interested in the effects of genotype, developmental temperature, selection regime, replicate population, and the interaction between selection regime and temperature. Note that because we are assaying recombination in heterozygous females (see Materials and Methods), we can only detect dominant genetic effects. Consistent with expectation, temperature significantly affects recombination fraction (*P* = 0.02, χ^2^ test; Figure 1, Table 2). In all three experimental evolution regimes, the proportion of offspring produced that is recombinant is higher at 25°C than it is at 16°C, though this increase is not statistically significant (*P* > 0.13, all comparisons, Wilcoxon Rank Sum Test). Thus, phenotypic plasticity in recombination fraction associated with temperature is observed across all three selective environments, though the magnitude of the effect is small.

Although the capacity for phenotypic plasticity is observed consistently across the three experimental evolution scenarios, these scenarios have yielded genetic divergence in recombination fraction independent of near-term rearing temperature. Specifically, the selective regime significantly contributes to the observed variation in recombination fraction among lines (*P* < 0.0001, χ^2^ test; Figure 1, Table 2). The H lines show the highest recombination fraction at both temperatures, the C lines show the lowest recombination faction at both temperatures, and the T lines show intermediate values of recombination fraction at both temperatures. For flies raised at 16°C, the recombination fraction of H lines is significantly higher than that of C lines (*P* < 0.0001, Tukey’s HSD). For flies reared at 25°C, the recombination faction of both H and T lines is significantly increased relative to the C lines (*P* < 0.03, both comparisons, Tukey’s HSD).

**Figure 1:**
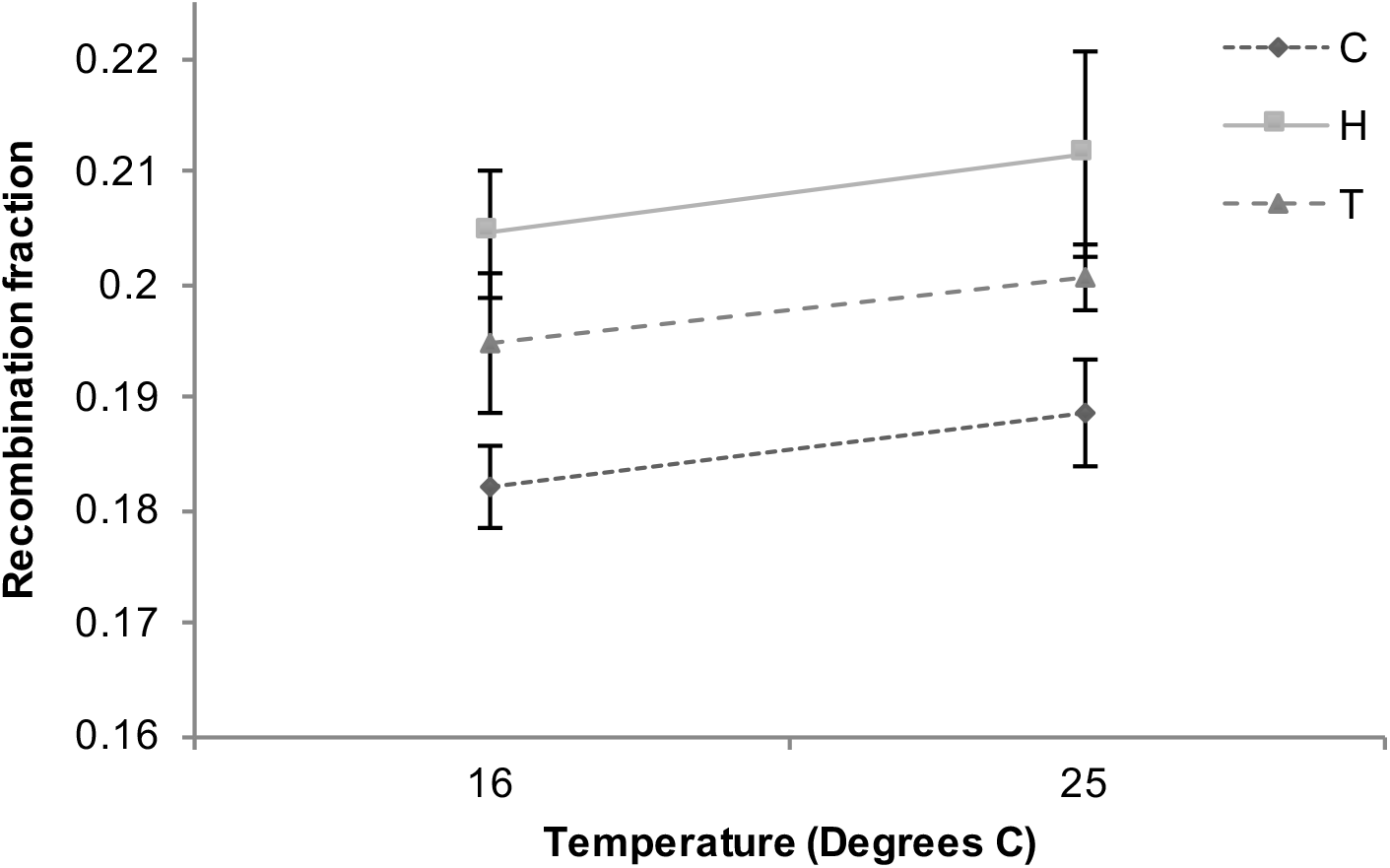
Average recombination fraction for each of the experimental evolution regimes at 16°C and 25°C. To obtain the regime average, the mean recombination fraction for each line was determined by averaging across replicates. The mean population recombination fraction was estimated by averaging across the average estimates for all of the lines in that population. The mean regime recombination fraction was estimated as the average across the five replicate populations. The standard error of the estimate across replicate populations is also shown for each regime at each temperature.

**Table 2:** Effect Tests for Logistic Regression on Recombination Fraction

| Source | df | $\chi^2$ | Prob > $\chi^2$ |
| --- | --- | --- | --- |
| Line | 145 | 1226 | **< 0.0001** |
| Regime | 2 | 111 | **< 0.0001** |
| Population | 12 | 149 | **< 0.0001** |
| Temperature | 1 | 5.4 | **0.020** |
| Regime x Temperature | 2 | 0.21 | 0.90 |

Other factors in the model that significantly contribute to the observed variation in recombination rate are genotype and population (*P* < 0.0001, both factors χ^2^ test; Figure 1, Table 2). This indicates that genetic differences among lines, even within populations and selective regimes, also contribute to phenotypic variation in recombination fraction. A significant effect of population indicates that replicates differ in their responses to the selective environment. This could be driven by random genetic drift over the course of the experimental evolution, or differences in the genetic variants present among replicates at their founding.

Experimentally-evolved populations do not differ in their degree of phenotypic plasticity in recombination fraction in response to temperature. That is, we find no significant interaction effect between ‘regime’ and temperature (*P* =0.90, χ^2^ test; Figure 1, Table 2). This indicates that there is no significant differentiation among selective treatments with respect to how recombination fraction changes in response to temperature. Indeed, the magnitude of the change in recombination fraction between 25°C and 16°C is consistent across the selective treatments (Figure 2).

**Figure 2:**
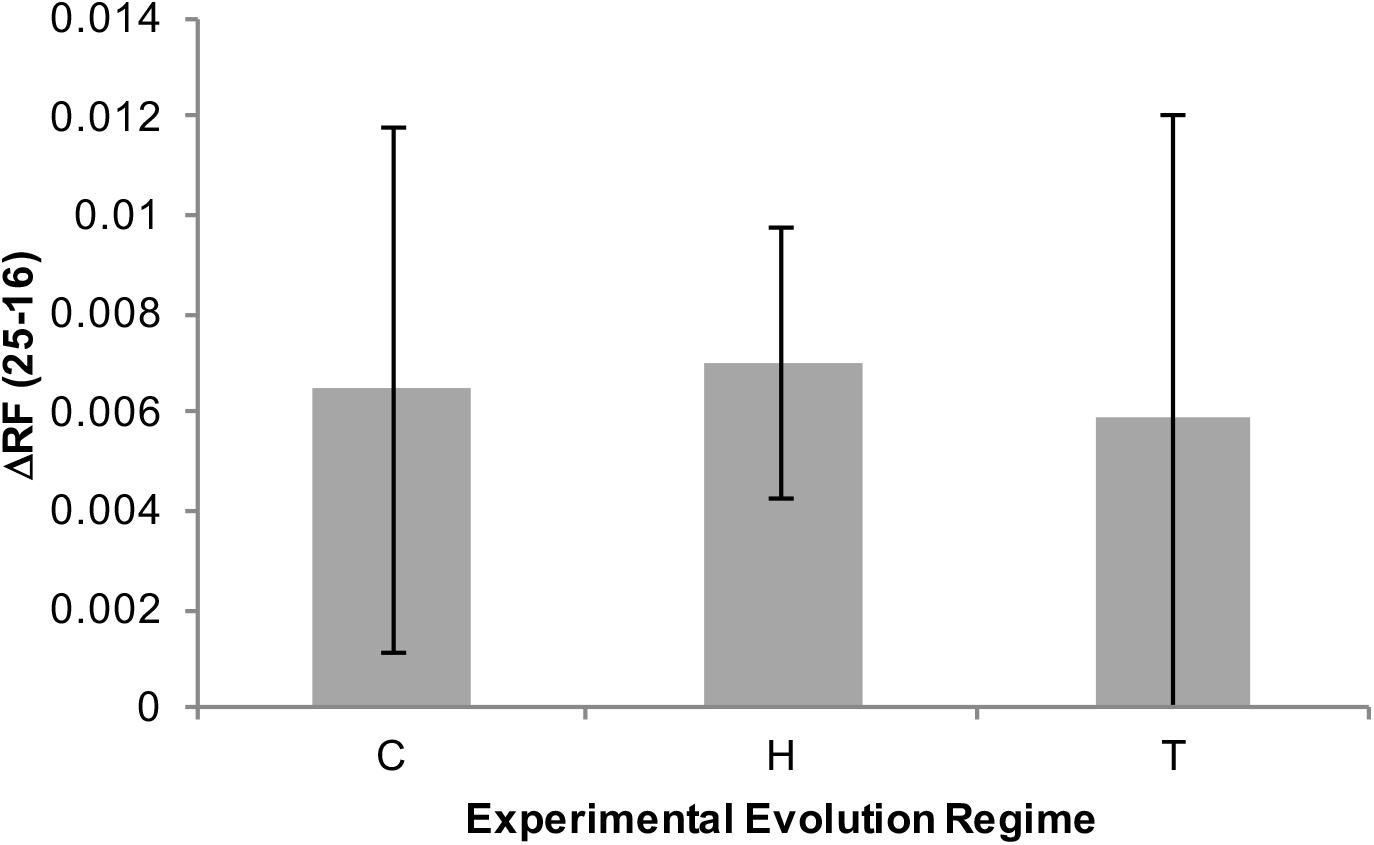
The average change in recombination fraction (ΔRF) between 25°C and 16°C for each experimental regime. The average change for each treatment was calculated as follows. The mean change for each line was estimated as the difference between the average recombination fraction across replicates at 25°C and the average recombination fraction across replicates at 16°C. The mean change for each population was then estimated as the average ΔRF across all lines in that population. Finally, we estimate average ΔRF for an evolution regime as the average across the five replicate populations. The standard error of that estimate across replicate populations is also shown for each regime.

To test whether there was genetic variation for recombination plasticity among lines, we also fit a model in which the response variable was the difference in recombination fraction between 25°C and 16°C. Our model indicates that there are no significant differences in the degree of plastic recombination among lines, populations, or experimental evolution regimes (*P* > 0.16, all comparisons; Table 3).

**Table 3:** Effect Tests for Model Fitting of Change in Recombination Fraction between 25°C and 16°C

| Source | df | Sum of Squares | F Ratio | Prob > F |
| --- | --- | --- | --- | --- |
| Line | 144 | 0.69 | 1.14 | 0.16 |
| Regime | 2 | 0.009 | 0.54 | 0.29 |
| Population | 12 | 0.06 | 1.19 | 0.90 |

## Discussion

### Temperature-associated plastic recombination

Phenotypic plasticity in recombination rate has been observed in a variety of taxa. Temperature in particular has been shown to affect the frequency of recombination in several species including Drosophila (Plough 1917, 1921; Smith 1936; Grell 1978), yeast (Johnston and Mortimer 1967), worms (Rose and Baillie 1979) and fungi (McNelly-Ingles et al. 1966; Rifaat 1969; Lu 1974). Our results confirm phenotypic plasticity in recombination fraction in response to temperature in *D. melanogaster*. It has been previously shown that recombination increases when flies are raised at temperatures higher or lower than their optimal temperature (Plough 1917, 1921; Smith 1936; Grell 1978). However, our data indicate that recombination fraction is lower at 16°C than it is than 25°C independent of the selective environment in which the flies were evolved. If one imagines that each population adapted to the temperature at which it was raised during the experimental evolution experiment, and departures from that optimal temperature would increase recombination as was seen before (Plough 1917, 1921; Smith 1936; Grell 1978), then one might have expected that in our study we would have found that the H lines would have higher recombination at 16°C than at 25°C, and vice-versa for the C lines. The overall reduction in recombination frequency at the lower versus the higher temperature in the current experiment is instead reminiscent of what is seen in *C. elegans*, where recombination frequency directly scales with temperature (Rose and Baillie 1979). A reduction in crossover frequency with decreased temperature has also been seen in yeast and Neurospora (Rifaat 1969), though Neurospora also shows evidence of increased recombination at lower temperatures (e.g. McNelly-Ingles et al. 1966).

At least two explanations for the varied recombinational responses to temperature within species and across experiments can be offered. First, genetic background clearly mediates recombination fraction, and different strains have been utilized across experiments. Genetic variation for recombination rate is clear not only in *D. melanogaster* (Broadhead et al. 1977; Brooks and Marks 1986; Hunter and Singh 2014; Hunter et al. 2016a), but also in many other species including mice, sheep, humans, and worms (e.g. Dumont et al. 2009; Rockman and Kruglyak 2009; Kong et al. 2010; Johnston et al. 2016). Differences in the genotypes of the strains used for experimentation may yield variable responses to temperature. Indeed, genotype-environment interactions significantly contribute to recombination rate variation in *D. melanogaster*, for instance (Hunter et al. 2016b). Second, the magnitude and direction of temperature-associated plastic recombination may vary across the genome. This is clearly the case in Drosophila, where centromeric regions show an exaggerated response to temperature (Plough 1921; Stern 1926), for example. The effect of temperature on recombination frequency is also heterogeneous across the yeast genome, but not in an obvious association with centromeres (Johnston and Mortimer 1967). Therefore, the differences among studies with regard to temperature-associated plastic recombination could be driven in part by different intervals of the genome being surveyed.

Why might recombination rates be sensitive to temperature? One possibility, as described above, is that populations are adapted to specific temperatures and being reared outside of these temperatures is stressful. Stress has long been associated with changes in recombination frequency (for review see Parsons 1988; Modliszewski and Copenhaver 2017). Another possibility is that recombination rates vary in response to temperature because the recombinational machinery is thermosensitive (Morgan et al. 2017). Specifically, if the proteins involved in the synaptonemal complex and axis formation function differently at different temperatures, then crossover number may vary according to temperature (Morgan et al. 2017). Note that this hypothesis is not at odds with the stress-associated recombination hypothesis; selection may shape thermotolerance of meiotic proteins directly or indirectly, and environments outside of the range to which individuals have adapted may lead to meiotic dysfunction (Morgan et al. 2017). If thermotolerance plays a role in temperature-associated recombination, then we might expect to see different thermostability of the meiotic axis and/or synaptonemal complex in our C versus H lines, as they have evolved distinct recombination rates.

### Genetic variation in recombination

It is well-documented that there is a genetic component to intraspecific variation in recombination rate. Such variation can be observed in humans, other mammals, plants, and insects (Shaw 1972; Valentin 1973; Dewees 1975; Hadad et al. 1996; Dumont et al. 2009; Johnston et al. 2016). Genetic variability in and heritability of recombination rate in Drosophila in particular has strong support in the literature. As noted above, classical genetic experiments indicate that the amount of crossing-over can vary among lines of D. *melanogaster* (Broadhead et al. 1977; Brooks and Marks 1986), even within a single population (Hunter et al. 2016a). It is therefore consistent with expectation that our analysis reveals that phenotypic variation in recombination fraction is explained in part by differences in genotypes (‘line’, Table 2).

Given that fluctuating environments favor recombination in certain circumstances, one initial hypothesis was that the variable temperature experimental evolution lines would evolve a higher baseline recombination rate. This was not observed, though our results do indicate divergence in recombination rate among the three experimental evolution regimes. That the T lines did not evolve higher recombination could indicate that although the environment varied cyclically with period two, the sign of linkage disequilibrium and/or epistasis did not change with the environmental changes.

The significant contribution of selective environment to the observed variation in recombination fraction in the current experiment suggests that recombination fraction was subject to different selective pressures in the three different environments. As a response to these pressures, the H lines evolved (or maintained) a higher recombination fraction independent of the temperature at which recombination was measured. C lines, in contrast, evolved a lower recombination fraction, which manifests at both temperatures. In contrast to our expectation, the T lines exhibit an intermediate phenotype at both temperatures. That the C lines have a lower recombination fraction than the H lines at both temperatures and recombination fractions at 16°C are consistently lower than recombination fractions at 25°C for all lines clearly indicates a role for temperature in both plastic recombination and baseline recombination rate. It should be noted that the map distance between the visible markers *ebony* and *rough* is 20.4 cM (Lindsley and Grell 1967), and the average distance between these markers in 112 lines from a North American population of *D. melanogaster* is 20.7 cM (Hunter et al. 2016a). These estimates are similar to the average recombination fraction of the H lines at 25°C (20.6 cM, Figure 1). This may indicate that the H lines maintained their recombination fraction while the C and T lines evolved a reduced recombination rate, but this is purely speculative. Were the founding population of these experimental evolution populations still available, this could be tested empirically.

Our data clearly indicate that baseline recombination rate evolves in response to temperature. We are as yet unaware of any data indicating clinal variation in recombination frequency among populations of any species, though our data suggest that there may be temperature-associated variation in this trait. Importantly, the adaptive significance of the evolved response to temperature observed in the current study remains unknown. Moreover, it is unknown whether changes in recombination among experimental treatments result from direct selection on recombination frequency itself or as an indirect consequence of selection on other traits. Both increases and decreases in recombination rate have been observed in laboratory selection experiments in which recombination rate itself was successfully subjected to artificial selection (Chinnici 1971; Kidwell 1972; Charlesworth and Charlesworth 1985). However, changes in recombination rate have been also shown to evolve as a correlated response to artificial selection on other characters (Flexon and Rodell 1982; Zhuchenko et al. 1985; Korol and Iliadi 1994; Rodell et al. 2004). It may be that the evolved changes in recombination rate among selection regimes are indirect consequences of natural selection on other phenotypes that are relevant for the experimental evolution treatments. Indeed, other studies on these lines have revealed divergence in other traits including body size, cell size, and metabolism across treatments (Adrian et al. 2016; Alton et al. 2016).

While selection appears to be driving recombination rate evolution among experimental evolution regimes, we cannot discount a role of random genetic drift in the evolution of recombination rate in these lines. Specifically, our data highlights variability in recombination rate that can be ascribed to replicate population. Thus, the phenotypic response to selection within a given treatment does vary among replicate populations. This variation could result from random genetic drift over the course of the experimental evolution course, or alternatively from stochastic variation in the pool of standing genetic variation present within each replicate at its founding. Previous work on these lines showed no effect of drift on the evolution of cell membrane plasticity among treatments (Cooper et al. 2012). This indicates that if our inter-replicate variability in the response to selection is indeed due to drift, then the strength of (direct or indirect) selection on recombination frequency is less intense than the strength of selection on cell membrane plasticity. Alternatively, if variance in recombination rate is driven by alleles of intermediate frequency, recombination rate could drift more rapidly than a trait driven by alleles of low frequency.

### Genetics of plastic recombination

Theory predicts that fluctuating environments can lead to the evolution of phenotypic plasticity under certain conditions. We thus hypothesized that the variable temperature experimental evolution lines would evolve a greater capacity for temperature-associated plastic recombination. In contrast to that expectation, here we show that while the capacity for plastic recombination is observed in all three experimental evolution treatments, the magnitude of temperature-associated plastic recombination is consistent across selection regimes. That is, there is no significant contribution of the interaction effect between the selection regime and temperature to observed phenotypic variation in recombination fraction.

While we observe no divergence among regimes in the capacity for plastic recombination, we note that divergence among regimes in phenotypic plasticity has been observed in other phenotypes. Specifically, T lines show an increased capacity for plasticity of the lipid composition of the cell membranes relative to the H and C lines (Cooper et al. 2012). These data illustrate that temporal variation in temperature can indeed lead to the evolution of increased plasticity in principle. That we see no evolution of an increased plasticity in recombination in lines subject to a variable thermal regime could suggest that there is little to no selective advantage of increased plastic recombination in environments that vary cyclically with respect to temperature Alternatively, it could be that the costs of greater plasticity in recombination are sufficiently large as to not be outweighed by the potential benefits of enhanced plastic recombination in variable environments.

When coupled with our observation that the selection treatments did yield divergence in baseline recombination frequency, our data indicate that the recombination fraction and temperature-associated plastic recombination have separable genetic architectures. This bears directly on a long-standing question on the genetic and molecular underpinnings of phenotypic plasticity. In particular, the extent to which genes underlying individual traits are the same genes underlying phenotypic plasticity in those traits remains controversial. Our data indicate that the genetic bases of these traits are at least partially non-overlapping in the case of recombination fraction in *D. melanogaster* and its response to temperature.

## Acknowledgments

The authors gratefully acknowledge Brandon Cooper for providing us with the lines that were used in the study. Brandon Cooper and Kristi Montooth provided valuable feedback on the experimental design. The authors also thank Brandon Cooper and Mohamed Noor for their valuable feedback on and suggestions for the manuscript. Comments from the associate editor and two anonymous reviewers greatly improved this manuscript. This work was supported by National Science Foundation grant MCB-1412813 to N. D. S.

---

**Supplemental Figure 1:**
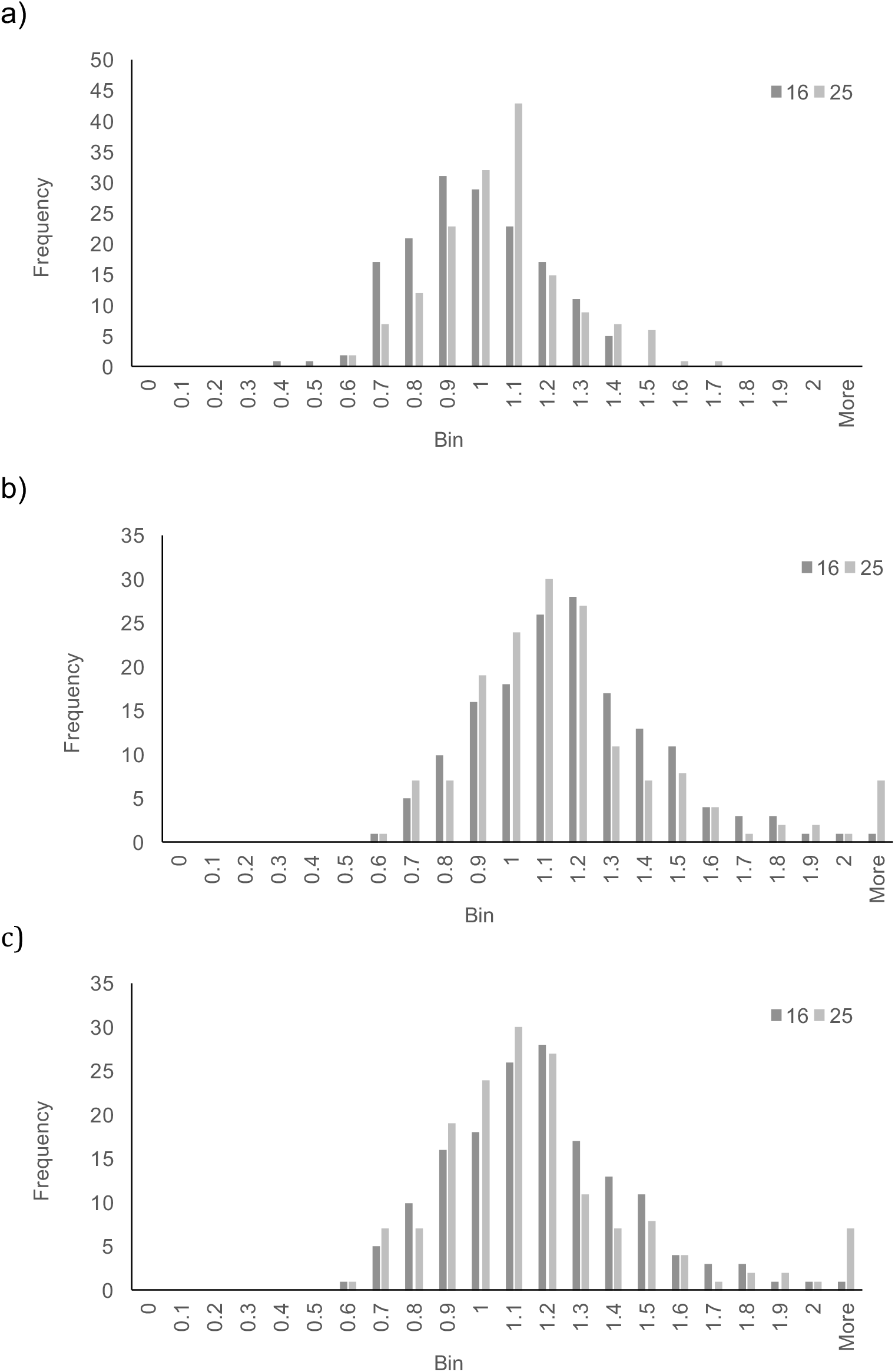
Distributions of ratios of a) males/females, b) wild-type/*e ro* flies, and c) *e* + / + *ro* flies at 16 degrees (dark grey) and 25 degrees (light grey) C.

